# Molecular Dynamics simulation of the E.*coli* FtsZ dimer

**DOI:** 10.1101/280230

**Authors:** Vidyalakshmi C Muthukumar

**Affiliations:** Department of Chemistry, University of Warwick

## Abstract

FtsZ dimer was studied to gain insights into FtsZ protofilament formation. In a previous simulation study of the *M.janaschii* dimer it was found that the monomer-monomer contacts in the GDP bound dimer is lower which results in the high curvature of the GDP bound protofilaments. In this study, we have simulated the *E.coli* FtsZ dimer. The initial structure was obtained from our previous study in which we had simulated the E.coli FtsZ monomer with its C-terminal IDR (Intrinsically Disordered region) built by homology modelling. The *M.janaschii* FtsZ dimer subunit contacts were used in the initial configuration. Simulations of the dimer were performed with GTP, GDP and ATP. We observed that the C-terminal domain rotates considerably during dimerization. We also observed the different dynamics of the GTP, GDP and ATP bound dimers due to which assembly into straight protofilaments is favoured only in the presence of GTP.

## 1. Introduction

FtsZ dimer structures have been crystallized from *M.janaschii* [1]. *In vitro*, the GTP bound protofilament is straight while the GDP bound protofilament has higher curvature [2]. Previous simulation study of the *M.janaschii* dimer also supports the higher curvature of the GDP bound form [3].

Since both in vivo and in vitro, the presence of the long, flexible C-terminal IDR was found to be essential for FtsZ polymerization [4], in our previous study, we had simulated the *E.coli* FtsZ monomer with GTP, GDP and without nucleotide. The simulations were performed on a homology model and included the C-terminal IDR. We had observed that the flexibility of this region depends on the nucleotide binding state of the monomer. A more flexible IDR was observed in the GDP bound monomer. We observed that GTP cannot bind to the monomer in a stable manner, its binding was variable, usually with the long chain amino acids of the S5- H4 loop. The central helix bends towards the C-terminal domain, this interaction disfavours stable binding with GTP (crystal structures reveal that interactions with the central helix are also responsible for holding the nucleotide in the binding site [5]). Some important conclusions from our previous study are 1) FtsZ has high intrinsic flexibility. The residues of the nucleotide binding region (HL1, S3 – H2A loop, S5 – H4 loop, the helix HL2, HL2 – H5 loop and the N-terminal residues of the central helix) have high flexibility. 2) The C-terminal domain showed small directed rotation in the direction of the nucleotide binding site in GTP bound state and away in the nucleotide free form. 3) The HL2-H5 recognizes the nucleotide binding state of the protein.

After studying the monomer interactions with the nucleotides, in this study we simulated the dimer to understand the formation of FtsZ protofilaments. *E. coli* FtsZ wild-type dimer was simulated with GTP, GDP and ATP to study a) the effect of dimerization on GTP binding b) the difference in interaction between the two chains in the GTP and GDP bound forms. We also simulated the wild-type dimer with ATP to compare it with GTP/GDP bound simulations. Two independent repeat simulations were performed for each system. The simulations are denoted by ‘WT-FtsZ dimer-GTP/GDP/ATP (simulation/repeat no.)’.

## 2. Methods

### 2.1 Dimer structure

To generate a PDB file for WT FtsZ dimer, WT protein structure (obtained from our previous MD simulation of *E. coli* FtsZ monomer with GDP) was aligned with chains A and B of the *Methanocaldococcus jannaschii* FtsZ dimer structure, 1W5A. This alignment was performed using the Multiseq alignment tool in VMD [6, 7] (Figure 2.3).

The starting position of the GDP nucleotide was obtained from the FtsZ monomer simulation with GDP after equilibration (from a representative frame at 93.9 ns). GTP was aligned manually in VMD using the position of GDP for reference. During manual alignment in VMD care was taken to keep a similar orientation of the nucleotide molecule and approximately, the same location in the nucleotide binding site. For ATP, a reference position was not available, therefore, it was simulated from a random initial position placed close to the nucleotide binding site. For verification, simulations were run for 100 ns and atleast 2 independent simulations were performed.

### 2.2 Molecular dynamics simulations

Atomistic molecular dynamics simulations were performed using GROMACS version 4.6.5 [8]. Simulations were performed in the AMBER94 force field [9, 10]. AMBER parameters for nucleotides developed by Meagher et al. [11] was used. Protein molecule was centred in the cubic box, at a minimum distance of 1.5 nm from box edges. Solvent (water) molecules were added in the coordinate file. SPC-E (Single Point Charge Extended) water model configuration was used [12]. Mg2+ and Cl- ions were added to neutralize the simulation box and at a minimum concentration of up to 10 mM (any of which was higher). Energy minimization was performed using the steepest descent minimization algorithm until the maximum force on any atom in the system was less than 1000.0 kJ/mol/nm. A step size of 0.01 nm was used during energy minimization. A cut-off distance of 1 nm was used for generating the neighbour list and this list was updated at every step. Electrostatic forces were calculated using Particle-Mesh Ewald method (PME) [13]. A cut-off of 1.0 nm was used for calculating electrostatic and Van der Waals forces. Periodic boundary conditions were used. A short 100 ps NVT equilibration was performed. During equilibration, the protein molecule was restrained. Leap-frog integrator MD simulation algorithm [14] was implemented with a time-step of 2 fs. All bonds were constrained by the LINear Constraints Solver (LINCS) constraint-algorithm [15]. Neighbour list was updated every 10 fs. A distance cut-off of 1 nm was used for calculating both electrostatic and van der Waals forces. Electrostatic forces were calculated using PME method. Two groups *i.e.* protein and non-protein (solvent, ligand and ions) were coupled with the modified Berendsen thermostat [16] set at 300 K. The time constant for temperature coupling was set at 0.1 ps. Long range dispersive corrections were applied for both energy and pressure. Coordinates were saved either every 2 ps or every 5 ps (in .xtc compressed trajectory format). A short 100 ps NPT equilibration was performed similar to the NVT equilibration with Parrinello-Rahman barostat [17, 18], with time constant of 2 ps, was applied to maintain pressure at a constant value of 1 bar. 100 ns MD simulation in NPT ensemble was implemented.

## 3. Results and Discussion

### 3.1 MD visualization

The simulations were viewed in VMD. Representative structures from well equilibrated simulation are provided (all structures are obtained at 93.9 ns). The two protein chains of the dimer molecule are denoted by ‘chain 1’ and ‘chain 2’. The corresponding nucleotides are also denoted as GTP-1 and GTP-2 respectively. The nucleotides of chain 1, GTP-1/GDP-1 (the lower FtsZ chain in the Fig.) are present at the dimer interface which is formed by the N-terminal domain of chain 1 and the C-terminal domain of chain 2 of the dimer.

It may be observed that the conformation of nucleotide binding site residues 44 – 72 *i.e.* the residues in HL1 region, the short beta strand S3 and the residues between S3 and H2A, is quite variable. The nucleotide GTP is present very much within the nucleotide binding site, Fig. 3.1 (a – b), whereas in our previous monomer simulations, GTP was present outside and could only form non-specific hydrogen bonds with the S5-H4 loop residues. The orientation of the guanine may differ to some degree, for example, between the chains (1) and (2) in the WT-FtsZ dimer-GTP (1), Fig. 3.1 (a). In the Fig. 3.1 (c) and (d) (WT-FtsZ dimer-GDP (1) and WT-FtsZ dimer-GDP (2)), it is observed that GDP is also present within the nucleotide binding sites of both the chains. In the Fig. 3.1 (e) and (f) (WT-FtsZ dimer-ATP (1) and WT-FtsZ dimer- ATP (2)), it is observed that ATP is present within the nucleotide binding site in the chain 2 and outside of the nucleotide binding site in the chain 1, in the simulation WT-FtsZ dimer-ATP (1). In the simulation WT-FtsZ dimer-ATP (2), the ATP molecule is present outside the nucleotide binding site in both the chains 1 and 2. ATP binding to the dimer appears to be variable/non-specific.

**Fig 3.1.**
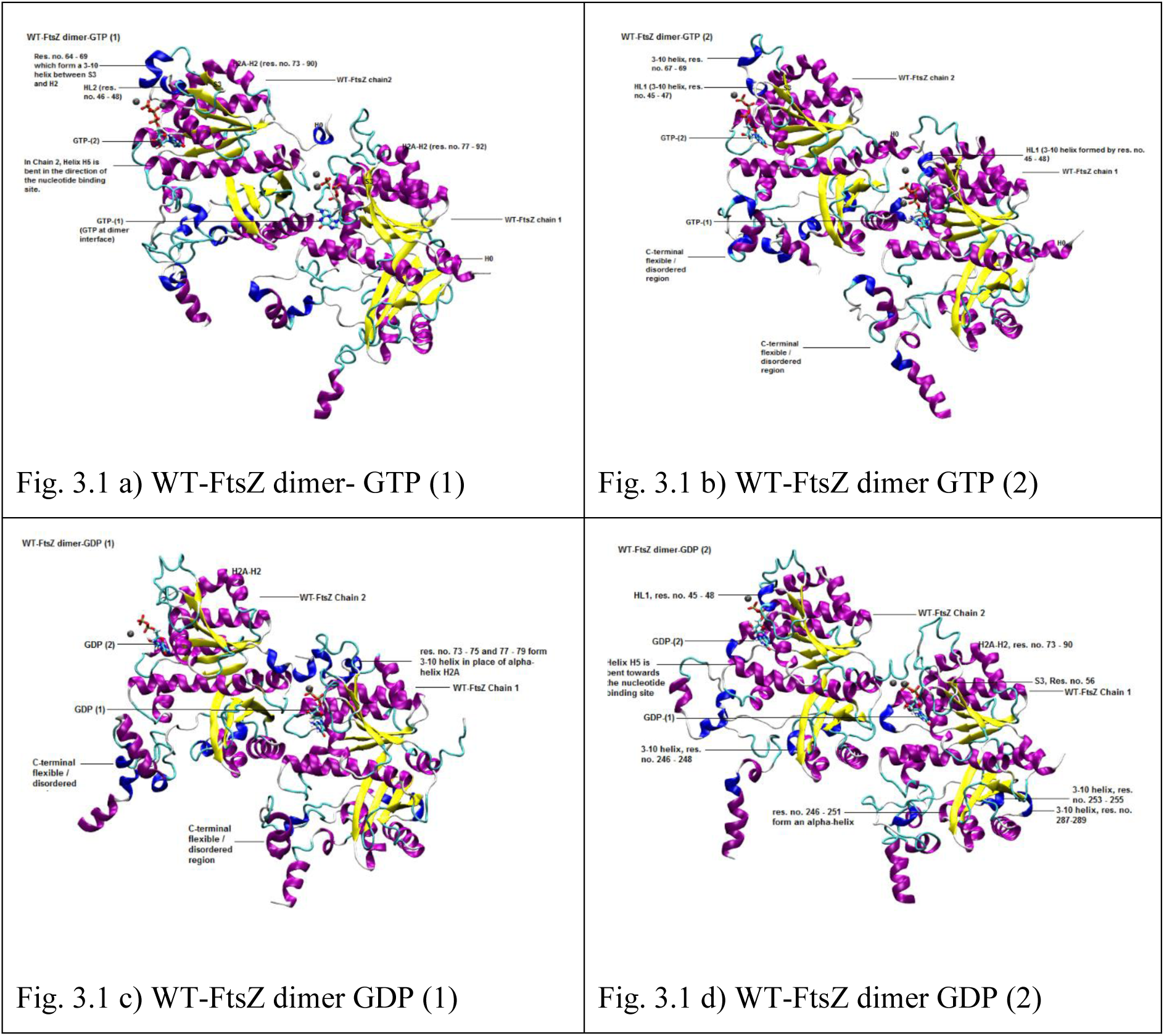
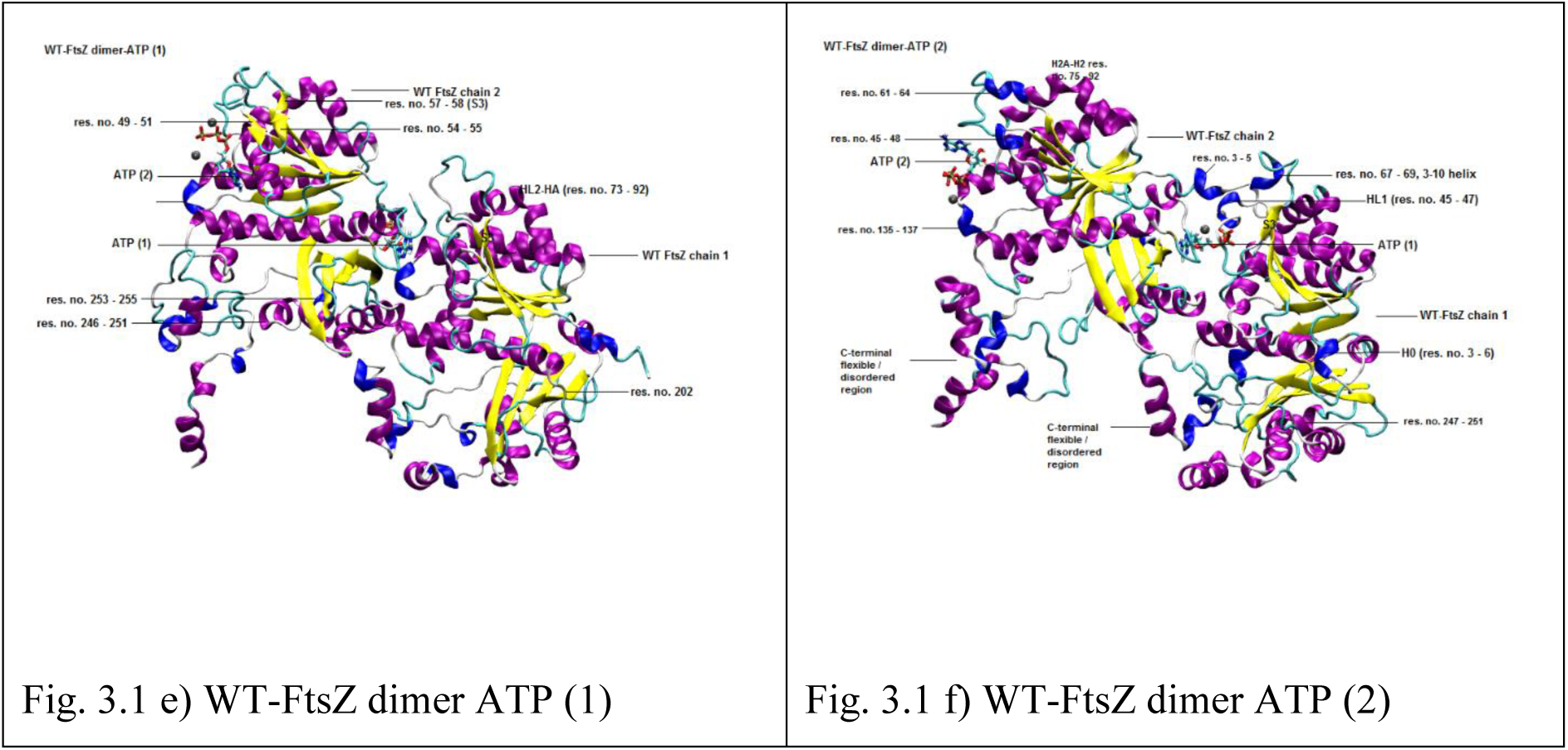
A-F): Structure of the proteins at 93.9 ns of molecular dynamics simulation. The structures of the protein are projected from the same angle in all the figures a - f. The molecules are coloured using the ‘structure’ colouring method in VMD. Mg^2+^ ions present within 5 nm of the nucleotides are shown in the figures. 1 – 2 Mg^2+^ ions are present.

In our previous study, we observed that in the simulations of the wild-type monomer with GTP, the central helix bends towards the C-terminal domain and away from the nucleotide binding site. Thereby, facilitating polymerization of the wild-type monomer in the presence of GTP (not with GDP). However, in the dimer simulations the central helix does not bend. This implies that the bending of the central helix observed in our previous simulations, is a feature of the FtsZ monomer binding with GTP. After formation of the dimer, the central helix does not bend towards the C-terminal domain, consistent with the expected subunit contacts formed between the N-terminal residues of the central helix of one subunit and HC3 of the other.

Conformational variability in the HC2-SC2 loop (245 – 257) of the C-terminal domain may be observed. For e.g., in the simulation, WT-FtsZ dimer-GTP (1), in chain 1, one 3-10 helix is observed (formed by residues 248 – 250), but in chain 2, two 3-10 helices are observed (formed by residues 246 – 249 and 253 – 255). In the simulation, WT-FtsZ dimer-GDP (1), in chain 1, a second α-helix is observed after HC2 (formed by residues 246 – 251).

It may be observed that the residues of the helix H0 and C-terminal IDR has different structures in the two chains and in all the simulations consistent with their high intrinsic disorder.

### 3.2 Root mean square deviation (RMSD)

RMSD was calculated for backbone atoms of the residues 12 – 311. Corresponding NPT equilibrated structures were used for reference structure. Protein coordinates were fitted on protein backbone atoms of residues 12 – 311 of chains 1 and 2. For most simulations, the RMSD profiles were stable (Fig. 3.2). RMSD fluctuations were high for two simulations, WT- FtsZ dimer-GDP (2) and WT-FtsZ dimer-ATP (2).

**Fig 3.2.**
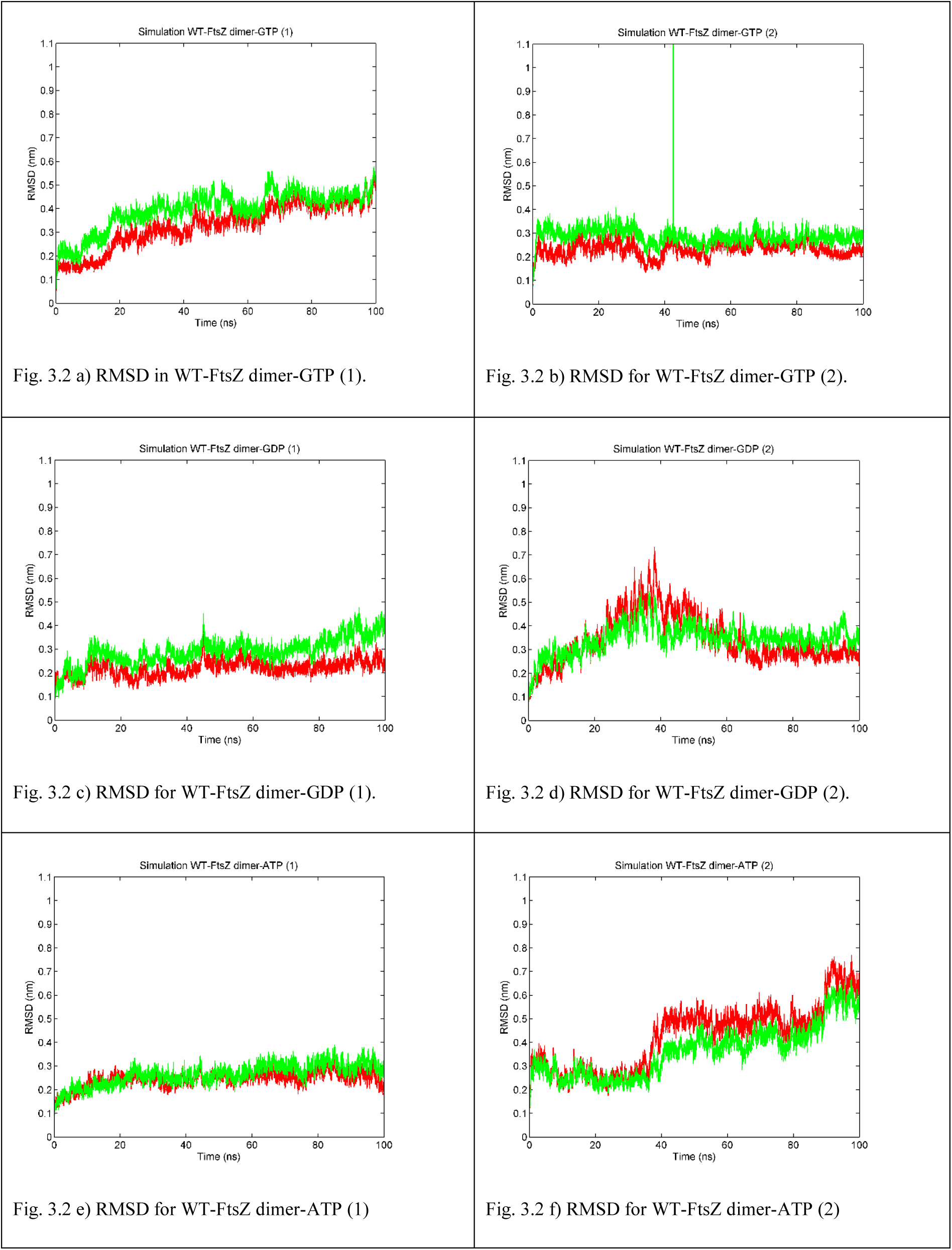
RMSD for simulations of wild-type FtsZ dimer with GTP, GDP and ATP. RMSD was calculated for backbone atoms of residues 12 – 311 of chain 1 (red) and chain 2 (green). RMSD for the six simulations of the wild-type FtsZ dimer are given in Fig. 7.2 a – f: a) WT-FtsZ dimer-GTP (1) b) WT-FtsZ dimer-GTP (2) c) WT-FtsZ dimer-GDP (1) d) WT-FtsZ dimer-GDP (1) e) WT-FtsZ dimer- ATP (1) f) WT-FtsZ dimer-ATP (2)

### 3.3 Root Mean Square fluctuation (RMSF)

RMSF is the standard deviation of atomic positions. It is a measure of flexibility of an atom during simulation. RMSF of protein backbone atoms (averaged per residue) was calculated. RMSF was calculated from 20 – 100 ns *i.e.* from the well equilibrated system. RMSF for the simulations are presented in Fig. 3.3.

**Fig 3.3.**
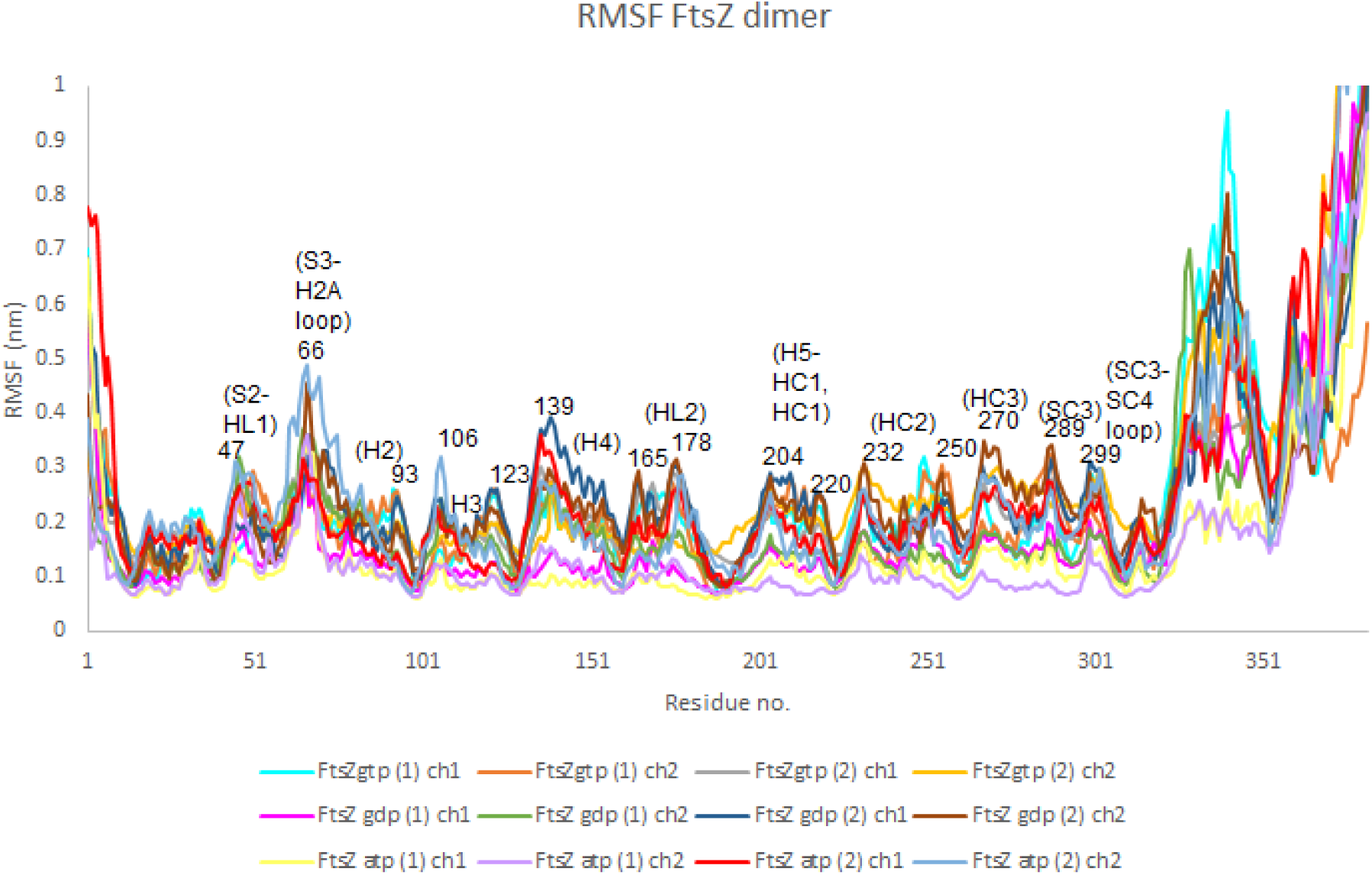
RMSF calculated for backbone atoms and averaged per residue, WT-FtsZ dimer-GTP (1) (chain 1 (cyan), chain 2 (orange)); WT-FtsZ dimer-GTP (2) chain 1 (grey), chain 2 (light orange); WT- FtsZ dimer-GDP (1) chain 1 (pink), chain 2 (green); WT-FtsZ dimer-GDP (2) chain 1 (dark blue), chain 2 (brown); WT-FtsZ dimer-ATP (1) chain 1 (yellow), chain 2(purple); WT-FtsZ dimer-ATP (2) chain 1 (red), chain 2 (light blue).

In our previous study of the wild-type monomer simulations (SI Fig. 1) higher RMSF (> 0.2 nm) was observed for HL1, S3-H2A loop, S5-H4 loop, HL2-H5, HC2 and HC3. In HC2 and HC3, sharp peaks were observed in which residues in the loops towards the end of helices or strands had higher RMSF. However, in the simulations of the wild-type dimer, there are two distinct regions in the RMSF plot. One is the high RMSF region of N-terminal residues 47 to 181 and the second is the high RMSF region of C-terminal residues 201 to 300. These two are separated by the low RMSF valley between 181 and 201 which corresponds to the central helix. Such an RMSF profile, indicates that the two domains rotate independently about the central helix. RMSF for residues 81 – 121 (which correspond to residues in H2A, H2, S4, H3) and residues 201 – 221 (which correspond to residues in H5-HC1 loop and HC1) have a much higher RMSF compared to the FtsZ monomer simulations (in our previous study). Higher RMSF in these residues neighbouring the central helix in the N-terminal and C-terminal domain reinforces that domain rotation occurs during FtsZ polymerization.

#### 1. RMSF for the simulations: WT-FtsZ dimer-GTP (1) and WT-FtsZ dimer-GTP (2)

In Supplementary Information Fig. 2 RMSF plot for each simulation is provided separately. For the simulation WT-FtsZ dimer-GTP (1) (SI Fig. 2 (a), (b)) it may be observed that the RMSF of the helix HL2, the HL2 – H5 loop and the N-terminal residues of the central helix for the chain 2 (green) is much lower than the RMSF of the corresponding residues in the chain 1 (red). In contrast, higher fluctuation of the following C-terminal ordered domain residues (from res. no. 190 onwards) is observed. This could be due to larger rotation of the C-terminal domain of the top chain, chain 2, during dimer formation. In addition, the N-terminal residues of the bottom chain (which forms the dimer interface with the C-terminal domain of the top subunit) is higher than the corresponding residues of the top chain. Higher RMSF in the contacting domains is consistent with dimer interactions resulting in domain motions which may be expected to be maximum in the immediate domains.

#### 2. RMSF for the simulations: WT-FtsZ dimer-GDP (1) and WT-FtsZ dimer-GDP (2)

RMSF for the simulations WT-FtsZ dimer-GDP (1) and WT-FtsZ dimer-GDP (2), are presented in SI Fig. 2 (c) and (d) respectively. In SI Fig. 2 (c), the chain 1 residues 12 – 311 (shown in red) has low RMSF, similar to the wild-type monomer simulations (reported in our previous studies). The N-terminal ordered-residues in chain 2 have higher RMSF than the corresponding residues in chain 1. The RMSF profile of the ordered-residues for WT-FtsZ dimer-GDP (2) for both the chains is similar to RMSF profile of WT-FtsZ dimer-GTP (1) the wild-type dimer simulations with GTP (described above), indicating that dimer interaction occurs.

Lower RMSF of the C-terminal domains of both chains in the simulation WT-FtsZ dimer-GDP (1) indicates that large domain rotation does not occur. Lower domain motion/rotation may be verified from the superposing of average coordinates (Fig. 3.6), in which the GDP bound dimer subunits undergo a lower rotation during dimerization than the GTP bound dimer subunits. Since the C-terminal domains do not rotate, dimer interaction would be weaker (consistent with the low RMSF of the chain-1 N-terminal domain shown in red). This is consistent with the lower monomer-monomer contact observed in the *M.janaschii* FtsZ dimer simulations resulting in high curvature of filaments. *In vitro*, lower polymerization ability due to weaker subunit interaction, is consistent with the high curvature or disassembly of GDP bound FtsZ.

#### 3. RMSF for the simulations: WT-FtsZ dimer-ATP (1) and WT-FtsZ dimer-ATP (2)

RMSF for the simulations WT-FtsZ dimer-ATP (1) and WT-FtsZ dimer-ATP (2), are presented in SI Fig. 2 (e) and (f) respectively. In SI Fig. 2 (e), the chain 1 and chain 2 ordered-residues 12 – 311 have low RMSF similar to the wild-type monomer simulations suggesting that dimerization is not efficient (following the discussion in points 1 and 2 above). We also observe a much lower RMSF for the C-terminal IDR for both the chains in the simulation. This may suggest that in the presence of a low flexibility IDR, polymerization does not occur efficiently. In contrast, the RMSF profile of the ordered-residues for WT-FtsZ dimer-ATP (2) for both the chains is similar to RMSF profile of WT-FtsZ dimer-GTP or GDP described above indicating that dimerization occurs.

### 3.4 Hydrogen bonds formed between nucleotides and FtsZ wild-type dimer

The residues are denoted by the three letter amino acid code and the residue no. ‘Main’ and ‘Side’ in the residue names indicates that the donor/acceptor atom belongs to the amino acid main chain and to the amino acid side chain respectively. The GTP molecule is denoted as ‘GTP-chain 1’ or ‘GTP-chain2’ referring to the corresponding chains. For calculating hydrogen bonds distance cut-off of 3.5 Å and angle cut-off 25° was used.

#### a) GTP

The GTP molecule in chain 1 (the bottom chain) in the simulation WT-FtsZ dimer-GTP (1) forms hydrogen bonds with ARG141-Side, THR131-Side in chain 1. The occupancy of the above hydrogen bonds are 86 %, 38 % respectively. The GTP molecule in chain 2 (the top chain) forms hydrogen bonds with ARG141-Side, GLU137-Side, PHE134-Main, ASN23-Side. The occupancy of the above hydrogen bonds are 81 %, 96 %, 46 %, 43 % respectively.

The GTP molecule in chain 1 (the bottom chain) in the simulation WT-FtsZ dimer-GTP (2) forms hydrogen bonds with ARG141-Side, ASN185-Side, THR131-Side in chain 1. The occupancy of the above hydrogen bonds are 87 %, 79 % and 41 % respectively. The GTP molecule in chain 2 (the top chain) forms hydrogen bonds with GLU137-Side, ARG141-Side and ASN23-Side. The occupancy of the above hydrogen bonds are 96 %, 65 % and 40 % respectively.

#### b) GDP

The GDP molecule in chain 1 (the bottom chain) in the simulation WT-FtsZ dimer-GDP (1) forms hydrogen bonds with GLU137-Side, ARG141-Side, THR131-Side in chain 1. The occupancy of the above hydrogen bonds are 98 %, 97 % and 62 % respectively. The GDP molecule in chain 2 (the top chain) forms hydrogen bonds with ARG141-Side. The occupancy of the hydrogen bond is 79 %.

The GDP molecule in chain 1 (the bottom chain) in the simulation WT-FtsZ dimer-GDP (2) forms hydrogen bonds with GLU137-Side and ARG141-Side. The occupancy of the above hydrogen bonds are 96 % and 96 % respectively. The GDP molecule in chain 2 (the top chain) forms hydrogen bonds with GLU137-Side, ARG141-Side and THR131-Side. The occupancy of the hydrogen bonds are 88 %, 88 % and 59 % respectively.

#### c) ATP

The ATP molecule in chain 1 (the bottom chain) in the simulation WT-FtsZ dimer-ATP (1) forms hydrogen bonds with GLU218-Side (chain 2), LYS140-Side, THR290-Side (chain 2) and LYS139-Side. The occupancy of the above hydrogen bonds are 93 % and 68 %, 60 % and 44 % respectively. The ATP molecule in chain 2 (the top chain) forms hydrogen bonds with ARG141-Side. The occupancy of the hydrogen bond is 85 %.

The ATP molecule in chain 1 (the bottom chain) in the simulation WT-FtsZ dimer-ATP (2) forms hydrogen bonds with ARG141-Side, ILE293-Main (chain 2), ILE293-Main (chain 2), VAL291-Main (chain 2). The occupancy of the above hydrogen bonds are 84 % and 61 %, 44 % and 38 % respectively. The ATP molecule in chain 2 (the top chain) forms hydrogen bonds with ARG141-Side and LYS140-Side. The occupancy of the hydrogen bond are 81 % and 48 % respectively.

Hydrogen bond analysis shows that GTP is bound in a much more stable manner than GDP in the dimer form (and in comparison to the GTP bound to the monomer in which variable/non-specific hydrogen bonds are observed). Therefore, dimerization or polymerization appears to be important for FtsZ GTPase activity which requires stable binding of nucleotide. The weak binding with GDP is essential for nucleotide exchange.

### 3.5 Principal Component Analysis

Principal component motion along the first eigen vector for the simulations are shown in Fig. 3.4 (a – f). In Fig. 3.4 (a) and (b), principal component motion along the first eigen vector for the simulations WT-FtsZ dimer-GTP (1) and (2) respectively are shown. In Fig. 3.4 (a), the top chain (chain 2) rotates left to right and the bottom chain (chain 1) rotates from right to left. In Fig. 3.4 (b), the top chain (chain 2) rotates anti-clockwise (left to right) and the bottom chain (chain 1) rotates clockwise (from right to left). However, the direction of rotation of the top chains (chain 2) and the bottom chains are equivalent in both Fig.3.4 (a) and (b). Rotation of the top chain anti-clockwise or left to right, would lead to capture of GTP by the N-terminal domain of the top chain. Hydrogen bond analysis supports this, since more number of hydrogen bonds are formed in the top chain in both the simulations with GTP. Simultaneously the C-terminal domain of the top chain also rotates anti-clockwise or left to right. The bottom chain N and C-terminal domains rotates clockwise or right to left. Such a motion in the bottom chain which also binds to the C-terminal domain of the top chain would open the nucleotide binding site of the bottom chain allowing rotation of the C-terminal domain of the top chain in anti- clockwise direction. Therefore, the N-terminal domain of the bottom chain performs the important function of facilitating GTP binding and hydrolysis in the top chain. We assume that GTP hydrolysis can occur in the top chain only because the rotation of the N-terminal domain of the top chain favours strong binding with GTP.

**Fig 3.4.**
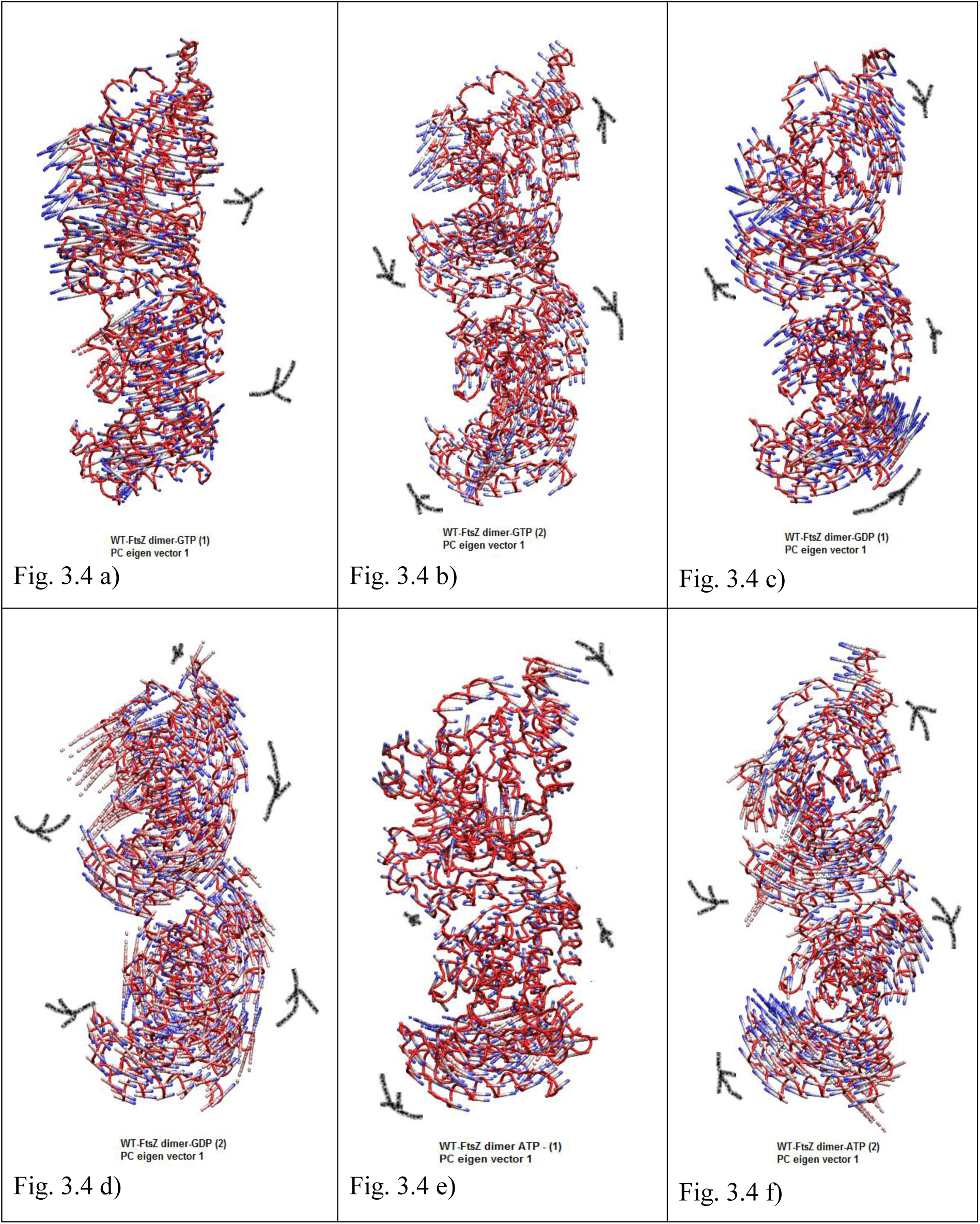
A-F) Principal component along eigen vector 1 for simulations of wild-type FtsZ dimer with nucleotides (a – b) GTP, (c – d) GDP and (e – f) ATP. PCA analysis was done for Ca atoms. The filtered trajectory along the first eigen vector is represented. The structure of the proteins are projected from the same angle in all states. The ‘tube’ representation in VMD is used to show initial frame (*i.e.* Frame 1, at 5 ps MD simulation). To show the motion of C-alpha atoms, the ‘CPK’ representation in VMD is used to show the frames from *t* = 0 to *t* = 100 ns with 1000 ps interval. VMD “trajectory” colouring method is used with the option ‘timestep’ (red is initial frame and blue final frame). The images were rendered in VMD. Arrows were drawn later to indicate the direction of rotation of N and C-terminal domains or the direction of rotation of the chains.

In Fig. 3.4 (c) and (d), principal component motion along the first eigen vector for the simulations WT-FtsZ dimer-GDP (1) and (2) respectively are shown. In contrast to the type of rotation of the wild-type dimer bound to GTP, the top and bottom chains rotate in the opposite direction of the corresponding chains in the GTP bound dimer form. In Fig. 3.4 (c) and (d), the top chains (chain 2) rotate clockwise and the bottom chains (chain 1) rotate anti-clockwise. The rotation of the N-terminal domain of the top chain in clockwise direction would open its nucleotide binding site and facilitate exchange or release of GDP. The rotation of the C-terminal domain of the top chain in clockwise direction would decrease the contact area between the two chains and thereby facilitate de-polymerisation. The bottom chain rotates anti- clockwise and again facilitates depolymerisation by decreasing the contact area with the top monomer by closing its nucleotide binding site.

In Fig. 3.4 (e) and (f), principal component motion along the first eigen vector for the simulations WT-FtsZ dimer-ATP (1) and (2) respectively are shown. In Fig. 3.4 (e), only the C-terminal domain of the bottom chain rotates anti-clockwise. In Fig 3.4 (f) the top and bottom chains rotate similar to top and bottom chains of the GTP bound dimers.

It may be noted here that the rotation observed in the C-terminal domain of the bottom chain is an independent domain motion of the C-terminal domain WT-FtsZ dimer-ATP (1) and not rotation of entire chain. Likewise, independent domain motions could be present in the other dimer simulations. Therefore, the observed rotation along the first eigen-vector is contributed by the independent domain motions (similar to what is observed in the simulation with ATP described above) and/or due to relative rotation of chains.

To distinguish between domain rotation and chain rotation, one of the domains only may be used for alignment. Therefore, the N-terminal domain ordered Cα atoms of the top monomer (residues 12 to 164 of the top monomer) was used. Average Cα coordinates were superposed for the simulations with GDP and with GTP. It was observed that, the bottom monomers were scattered (due to non-alignment) (Fig. 3.5 (a)). Therefore, considerable rotation of bottom chain about the top chain occurs in the dimer. Between the GTP and GDP forms of the dimer, it may be observed that the C-terminal domain of the top monomer is rotated or shifted in the simulations with GTP (Fig. 3.5 (b) to favour dimer interaction. The rotation of C-terminal domain of the bottom monomers may also be observed (3.5 (c)). Similar observation was made when the GTP and GDP bound FtsZ monomers from our previous study were compared (Fig. 3.5 (d)).

**Fig 3.5.**
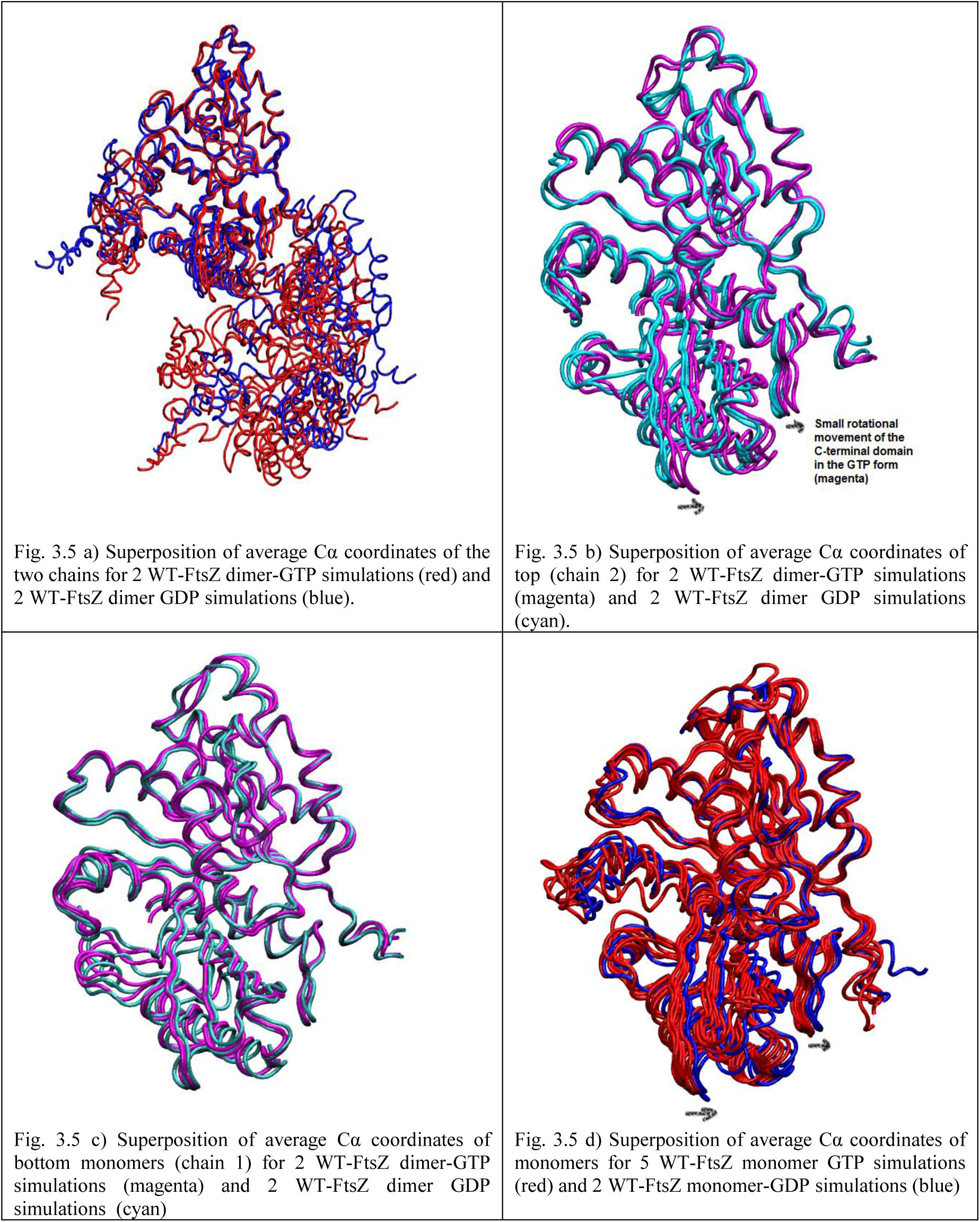
A – D) Superposition of average Ca coordinates of the top and bottom chains from different simulations with GTP and GDP. Alignment based on the N-terminal ordered residues (residue 12 – 164). (a) average Ca coordinates of the two chains for 2 WT-FtsZ dimer-GTP simulations (red) and 2 WT-FtsZ dimer GDP simulations (blue), (b) top (chain 2) for 2 WT-FtsZ dimer-GTP simulations (magenta) and 2 WT-FtsZ dimer GDP simulations (cyan), (c) bottom monomers (chain 1) for 2 WT-FtsZ dimer-GTP simulations (magenta) and 2 WT-FtsZ dimer GDP simulations cyan), (d) 5 WT-FtsZ monomer GTP simulations (red) and 2 WT-FtsZ monomer-GDP simulations (blue)

However, when the same comparison was done for the top or bottom monomers of the WT- FtsZ dimer-GTP simulations and the WT-FtsZ monomer simulations, it was observed that the C-terminal domain of the top monomer has rotated considerably Fig. 3.6 (a – b). When the GDP bound monomer and the dimer subunits are compared, same observation is made but the rotation is smaller than in the GTP dimer (Fig. 3.6 c – d). Therefore, it is clear that the subunits of the protofilament differ from the monomer by rotation of the C-terminal domain. This implies that domain rotation occurs during polymerization of FtsZ. It depends more on effective polymerization of FtsZ and less on the nucleotide binding state.

**Fig 3.6.**
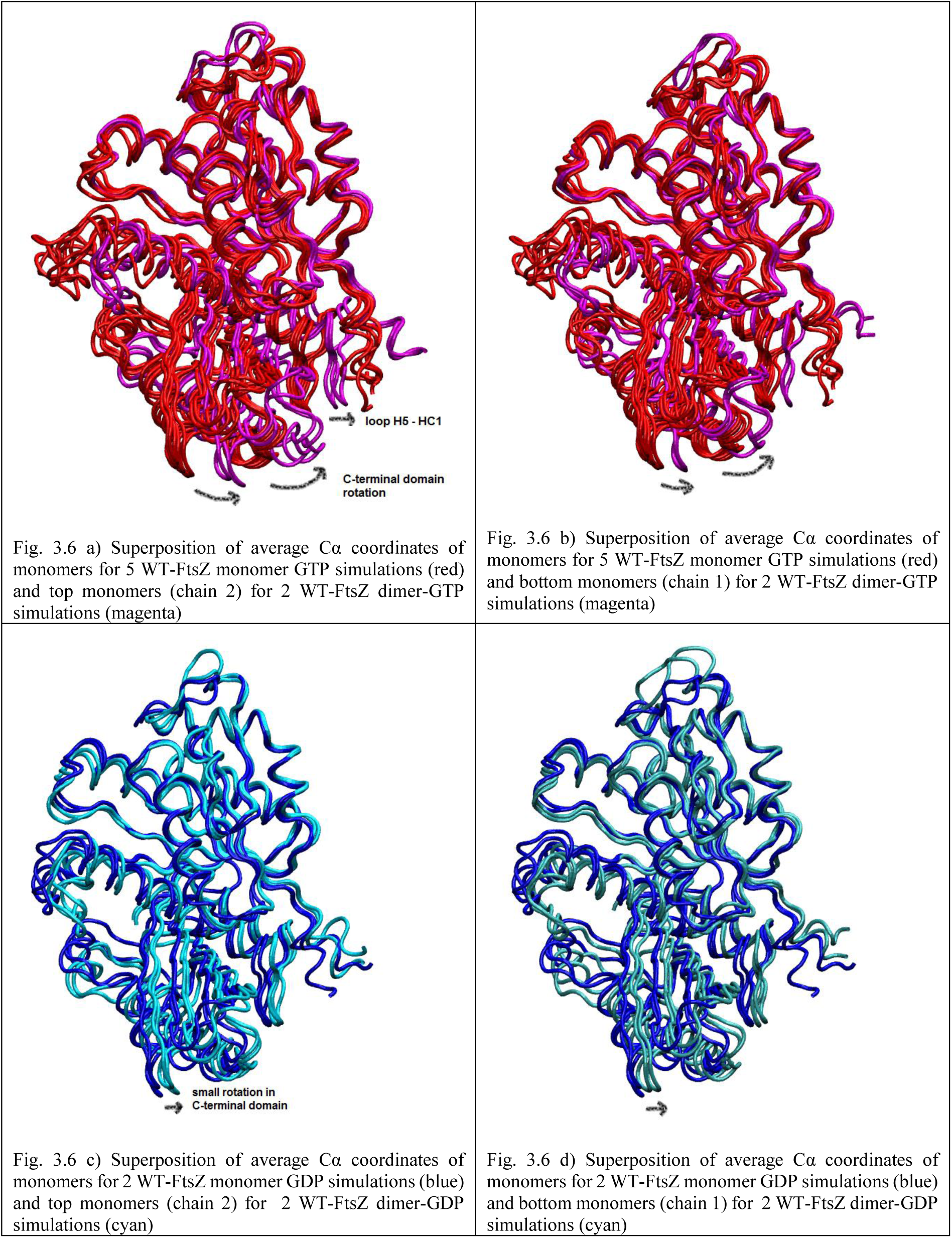
A – D) Superposition of average Ca coordinates of the top and bottom chains from different simulations with GTP and GDP. Alignment has been performed on the N-terminal ordered residues (residue 12 – 164). (a) 5 WT-FtsZ monomer GTP simulations (red) and top monomers (chain 2) for 2 WT-FtsZ dimer-GTP simulations (magenta), (b) 5 WT-FtsZ monomer GTP simulations (red) and bottom monomers (chain 1) for 2 WT-FtsZ dimer-GTP simulations (magenta), (c) 2 WT-FtsZ monomer GDP simulations (blue) and top monomers (chain 2) for 2 WT-FtsZ dimer-GDP simulations (magenta), (d) 2 WT- FtsZ monomer GDP simulations (blue) and bottom monomers (chain 1) for 2 WT-FtsZ dimer-GDP simulations (magenta)

In the crystal structure of *S.aureus* FtsZ [19] with GDP, it was observed that the crystal had the properties expected for the GTP bound FtsZ (with respect to the HL2 – H5 region, the T7 loop position inside the nucleotide binding site of the adjacent subunit). The investigators, consider it likely that the crystal was obtained soon after GTP hydrolysis, therefore, could have features of the GTP form. In their crystal structure, the C-terminal domain was rotated towards the N- terminal domain of the adjacent subunit. The degree of rotation was about 25°. We observe a rotation of about 10° along SC1 (line drawn between residues 223 and 228 in SC1) between the GTP bound monomer structures and the subunits of the GTP bound dimer. RMSD between FtsZ GTP monomer structures lie between 1.2 – 1.8 Å (alignment based on Ca atoms of residues 12 – 311). However, the RMSD between the GTP bound monomer and GTP bound top chain in the dimer forms are 4 – 4.5 Å. Therefore, our study supports the C-terminal domain rotation of FtsZ. We find that such conformational change occurs during dimerization or protofilament formation. It does not depend significantly on the nucleotide binding state of the protein.

## Conclusions

The *E. coli* FtsZ dimer was simulated with GTP, GDP and ATP. The structure of the dimer was built using the monomer structure from one of our previous simulations of the monomer with GDP. The nucleotides GDP and GTP were added in the simulations after approximate alignment with the position of GDP in the monomer simulation WT-FtsZ mono-GDP (1). The nucleotide ATP was added in the simulations at an approximate position in the nucleotide binding site. The C-terminal flexible / disordered region has high flexibility. GTP binds to the FtsZ dimer in a stable manner. The GTP molecule of the top chain forms hydrogen bonds with the residues of the S5-H4 loop (Arg141, Glu137, Phe134) and Asn23 (in the helix H1) in both the simulations. In contrast, the GTP molecule of the lower chain forms fewer hydrogen bonds, two hydrogen bonds in the simulation WT-FtsZ dimer-GTP (1), with Arg141 and Thr131; and three hydrogen bonds in the simulation WT-FtsZ dimer-GTP (2), with Arg141, Asn185 and Thr131. Hydrogen bonds with Asn23 of the helix H1 is not present. What happens in the presence of a third subunit, remains to be seen. On comparing the hydrogen bonds of the simulations of the wild-type dimer with GTP and the wild-type dimer with GDP, it may be observed that fewer hydrogen bonds are formed in the simulations of the wild-type dimer with GDP, for e.g. in the chain 2 of the dimer simulation WT-FtsZ dimer-GDP (1), only one hydrogen bond was formed with occupancy >38 % (with Arg141, occupancy 79%). This suggests the hydrogen bond interaction of the wild-type dimer is stronger with GTP than with GDP which is in accordance with the weaker binding of GDP into the binding site (as seen from measurements of exchange).

In contrast to GTP binding with the wild-type monomer which resulted in the opening of the nucleotide binding site by bending of the central helix H5 towards the C-terminal domain (reported in our previous simulations of the monomer), no such bending of the central helix was observed in the wild-type dimer simulations. From PCA analysis we observed that the rotation of the two subunits in the dimer was such that polymerization is facilitated in the GTP bound dimer form and depolymerisation is facilitated in the GDP bound form. Comparison of average structures from our previous monomer simulations with the subunits of the dimer revealed that C-terminal domain rotation occurs during FtsZ polymerization. There is a large rotation of the C-terminal domain in the subunits of the GTP bound dimer and smaller rotation in the GDP bound dimer. It appears that during the rotation of the C-terminal domain, the Asp300 of SC3-SC4 comes in contact with the Arg32 of H1-S2 and a clamp is formed by the two residues. With a clamped H1 helix, GTP is able to bind to Asn23 of the helix H1. In contrast in the bottom subunit the C-terminal domain rotation is lower, with the clamp not in place, GTP is unable to bind to Asn23. During this clamp formation, outer structure of the N-terminal domain are displaced (rotation like motion of the N-terminal domain is observed). Structural changes are transmitted through the clamp to N-terminal domain (S2, HL1, S3-H2A, H2, S4, H3, S5, H4).

In the simulation, WT-FtsZ dimer-ATP (2), it was observed that the two chains rotate according to the GTP bound dimer subunits. Therefore, it may be said that in the absence of a well bound nucleotide also, the chains may undergo rotation. Rotation occurs as a result of FtsZ dimerization. In the simulation WT-FtsZ dimer-ATP (1), it may be observed that the RMSF for the C-terminal IDR is much lower indicating that the flexibility of the C-terminal IDR affects subunit subunit interactions.

On viewing the filtered trajectory along the first eigen vector for the simulation WT-FtsZ dimer GTP (1), it appears that the IDR of the bottom chain expands in two directions, towards the HL2-H5 of the bottom subunit and the C-terminal domain of the other. Such a two-directional movement in the C-terminal IDR may be verified from the structures at 93.9 ns in Fig. 3.1 (a) bottom chain, 3.1 (d) top chain, 3.1 (f) bottom chain leading to right to left or clockwise rotation in respective chains. This leads to rotation of the bottom chain (right to left or clockwise) and also changes the domain orientation with respect to each other. Different rotational patterns (right to left or clockwise) observed in PCA analysis raises the possibility that the protofilament is highly dynamic and may adapt to several subunit configurations. This may allow a range of subunits per unit volume which may also be investigated using MD simulations. Thus permitting FtsZ assemblies from protofilaments with different curvatures, rings and sheets observed *in vitro* [20].

## Acknowledgement

We would like to thank the Marie Curie Actions program for funding the project, the Minerva supercomputer resource in Warwick University and Iridis supercomputer in Southampton University, Dr. Alison Rodger and Dr. Syma Khalid for supervision in the project.

